# Role of Atg1 in morphologic changes of the pathogenic fungus *Trichosporon asahii*

**DOI:** 10.1101/2025.06.16.659855

**Authors:** Mei Nakayama, Yasuhiko Matsumoto, Sanae Kurakado, Takashi Sugita

## Abstract

*Trichosporon asahii* is a dimorphic fungus that causes severe invasive fungal infections, particularly in patients with neutropenia. Depending on nutrient availability, *T. asahii* exists in yeast, hyphae, or arthroconidia forms. Autophagy, a cellular degradation pathway that removes old or damaged organelles, is essential for the survival of many eukaryotic organisms under nutrient-limited conditions. Atg1 is a key regulator of the early phases of autophagy, especially under nitrogen starvation. The role of Atg1 in regulating morphology, stress resistance, or virulence in *T. asahii*, however, remains poorly understood. Here, we generated three *atg1* gene-deficient *T. asahii* mutants and investigated their phenotypic characteristics to reveal the role of Atg1 in *T. asahii*. The *atg1* gene-deficient mutants exhibited no growth defects under high-temperature or various chemical stress conditions, including antifungal drugs. The mutants exhibited an increased proportion of hyphal cells when cultured in Sabouraud dextrose broth (SB), a medium commonly used for fungi. On the other hand, no morphologic differences were observed between the parent strain and the *atg1* gene-deficient mutants under a nitrogen-limited condition. The virulence of these *atg1* gene-deficient mutants was maintained in a silkworm infection model. Furthermore, all three generated *atg1* gene-deficient mutants exhibited consistent phenotypes. Our findings suggest that while Atg1 does not play a major role in stress tolerance or virulence in *T. asahii*, it plays a role in regulating its dimorphic morphologic changes.

## 1. Introduction

*Trichosporon asahii* is a dimorphic fungus that exists in yeast, hyphal, and arthroconidia forms (*1*). *T. asahii* causes severe invasive fungal infections, particularly in immunocompromised hosts, such as those with neutropenia (*2–4*). Infections caused by *T. asahii* can lead to severe deep-seated mycoses, including urinary tract and disseminated infections (*5*,*6*). Furthermore, *T. asahii* exhibits tolerance to echinocandin antifungal agents commonly used in clinical settings, often leading to breakthrough infections in patients receiving micafungin (*5*,*7*). The ability of *T. asahii* to form biofilms on medical device surfaces, such as catheters, further complicates treatment by conferring resistance to various antifungal drugs and contributing to chronic, treatment-resistant infections (*8*). These characteristics of *T. asahii* emphasize its clinical significance as a pathogenic fungus.

Autophagy is a conserved cellular process that maintains cellular homeostasis by degrading and recycling cytoplasmic components, including damaged organelles and misfolded proteins (*9*,*10*). Autophagy is regulated by signaling pathways such as mTOR and AMPK and is activated under stress conditions such as nutrient deprivation (*11*,*12*). In the model eukaryote *Saccharomyces cerevisiae*, autophagy plays a critical role in survival under nutrient-limited conditions (*13*,*14*). The process is initiated in response to nutrient deprivation, with Atg1 acting as a key regulator in its early stages (*9*,*15*). Atg1 and other autophagy-related proteins are essential for the progression of autophagy and are required for stress resistance and pathogenicity in *Cryptococcus neoformans*, a basidiomycete closely related to *T. asahii* (*16*,*17*). In *Candida albicans*, another dimorphic fungus, Atg1 is required for biofilm formation (*18*,*19*). The role of the autophagy-related protein Atg1 in morphologic transitions, stress resistance, and virulence in *T. asahii*, however, remains unclear.

In the present study, we independently generated three *atg1* gene-deficient mutants of *T. asahii* and evaluated their morphology, stress sensitivity, and virulence. Our results revealed that *atg1* gene deficiency promotes hyphal formation under culture conditions using Sabouraud dextrose broth (SB), a medium commonly used for fungi, but does not affect growth under various stress conditions or virulence in the silkworm infection model. These findings suggest that Atg1 is involved in regulating the morphologic changes of *T. asahii*.

## 2. Materials and Methods

### 2.1. *T. asahii* strains

*T. asahii* MPU129Δ*ku70* was used as the parental strain (*20*). Three independent *T. asahii atg1* gene-deficient mutants were generated in this study (Table 1).

**Table 1.** *T. asahii* strains used in this study.

| *T. asahii* strains | Relevant genotype | Background | Reference |
| --- | --- | --- | --- |
| MPU129 $\Delta ku70$ (Parent strain) | *ku70::nptII* | MPU129 | Matsumoto Y., *et. al.* (22) |
| $\Delta atg1$ #1-#3 | *ku70::nptII, atg1::NAT1* | MPU129 $\Delta ku70$ | This study |

### 2.2. Culture conditions of *T. asahii*

*T. asahii* strains were cultured according to previously described methods (*20*). *T. asahii* strains were grown on Sabouraud dextrose agar (SDA; 1% hipolypeptone, 4% dextrose, and 1.5% agar) containing G418 (100 µg/mL) and cefotaxime (100 µg/mL) at 27°C for 1–2 days.

### 2.3. Reagents

Nourseothricin was purchased from JENA BIOSCIENCE GMBH (Dortmund, Germany). Fluconazole was purchased from Tokyo Chemical Industry Co., Ltd. (Tokyo, Japan). Cefotaxime sodium, D-glucose, agar, NaCl, sorbitol, H O, dithiothreitol (DTT), and Yeast Nitrogen Base without Amino Acids and Ammonium Sulfate were obtained from FUJIFILM Wako Pure Chemical Corporation (Osaka, Japan). Congo red was purchased from Sigma-Aldrich Co., LLC (St. Louis, MO, USA). G418 was obtained from Enzo Life Science, Inc. (Farmingdale, NY, USA). Hipolypeptone was purchased from NIHON PHARMACEUTICAL Co., LTD. (Tokyo, Japan). Tunicamycin was purchased from Cayman Chemical Company (Ann Arbor, MI, USA). Amphotericin B and KCl were obtained from Wako Pure Chemical Industries, Ltd. (Osaka, Japan). Sodium dodecyl sulfate (SDS) was purchased from NIPPON GENE Co., LTD. (Tokyo, Japan). Tween 80 was obtained from MP Biomedicals LLC (Santa Ana, CA, USA).

### 2.4. Conservation of Atg1 protein amino acid sequence in *T. asahii* and other fungi

Homology and domain analyses were performed as previously described (*21*). The amino acid sequence of the *T. asahii* Atg1 protein was obtained from the National Center for Biotechnology Information (NCBI) (https://www.ncbi.nlm.nih.gov/). BLASTp (https://blast.ncbi.nlm.nih.gov/Blast.cgi) was used to compare the sequence homology with the Atg1 proteins of *C. albicans*, *A. fumigatus*, *C. neoformans*, and *S. cerevisiae*. A phylogenetic tree was constructed using the maximum likelihood method with MEGA11 Software (https://www.megasoftware.net/). The NCBI gene accession numbers of Atg1 proteins used for phylogenetic analysis and alignment are listed in Table 2. The NCBI domain accession numbers used for functional domain analysis and alignment are shown in Table 3.

**Table 2.** NCBI gene accession numbers for the Atg1 protein used to create the phylogenetic tree.

| Species | Definition | NCBI accession | Size<br>(aa) | Query<br>Cover<br>(%) | Percent<br>Identical<br>(%) |
| --- | --- | --- | --- | --- | --- |
| *Trichosporon asahii* | hypothetical protein A1Q1_02656<br>(*Trichosporon asahii* var. *asahii* CBS 2479) | XP_014179392.1 | 995 | 100 | 100 |
| *Cryptococcus neoformans* | ULK/ULK protein kinase<br>(*Cryptococcus neoformans* var. *grubii* H99) | XP_012048654.1 | 985 | 98 | 51.87 |
| *Aspergillus fumigatus* | serine/threonine protein kinase<br>(Pdd7p), putative (*Aspergillus fumigatus* Af293). | XP_751920.1 | 973 | 77 | 52.33 |
| *Candida albicans* | serine/threonine protein kinase<br>(*Candida albicans* SC5314) | XP_717245.2 | 834 | 31 | 50.46 |
| *Saccharomyces cerevisiae* | serine/threonine protein kinase ATG1<br>(*Saccharomyces cerevisiae* S288C) | NP_011335.1 | 897 | 61 | 47.45 |

**Table 3.**
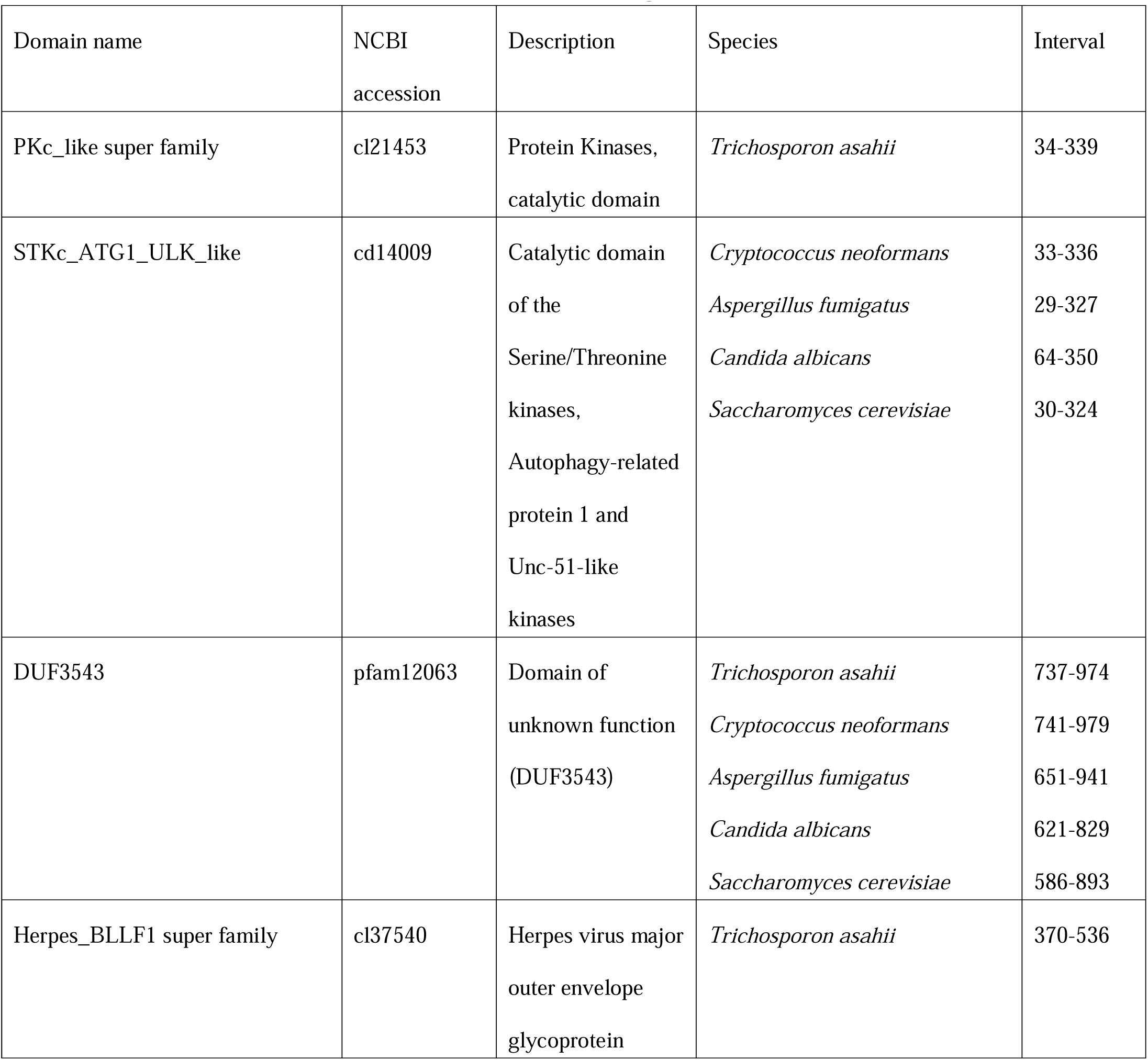
NCBI domain accession number for the Atg1 protein.

### 2.5. Generation of *atg1* gene-deficient *T. asahii* mutants

Generation of the gene-deficient *T. asahii* mutants was performed according to previously described methods (*22*). The pUC19-*atg1* (5’UTR) -*NAT1*-*atg1* (3’UTR) plasmid was constructed using the In-Fusion method. Approximately 1,200 bp of the 5′-UTR and 3′-UTR regions of the *atg1* gene were cloned upstream and downstream of the *NAT1* gene region in the pUC19-NAT1 vector using the In-Fusion method (In-Fusion Snap Assembly Master Mix; Takara Bio Inc., Shiga, Japan). Using the pUC19-*atg1* (5’UTR) -*NAT1*-*atg1* (3’UTR) plasmid, polymerase chain reaction (PCR) was performed to amplify a DNA fragment containing *atg1* (5’UTR) -*NAT1*-*atg1* (3’UTR). The primers used for PCR are listed in Table 4. Gene transfer to *T. asahii* cells was performed as described previously with modifications (*23*). The parental strain was cultured on SDA at 27°C for 1 day. Cells on the agar medium were suspended in 2 mL of saline and filtered through a 40-µm cell strainer. The filtered suspension was centrifuged at 10,000 rpm for 5 min, and the pellet was resuspended in 1 mL of distilled water and centrifuged again under the same conditions. This washing step was repeated four times. The washed cells were then suspended in 1 mL of 1.2 M sorbitol solution and centrifuged at 10,000 rpm for 5 min. The harvested pellet was resuspended in 40 µL of 1.2 M sorbitol to obtain competent cells. To perform electroporation, 4 µL of the PCR-amplified *atg1* (5’UTR)-*NAT1*-*atg1* (3’UTR) DNA fragment was added to 40 µL of the competent cells and incubated on ice for 15 min. The mixture was transferred to a 2-mm gap cuvette (Bio-Rad Laboratories, Inc., CA, USA), and electroporation was performed using a Gene Pulser Xcell (Bio-Rad Laboratories, Inc., CA, USA) with the time constant protocol (1800 V, 5 ms). After electroporation, 500 µL of Yeast Polypeptone Dextrose (YPD) medium (2% hipolypeptone, 1% yeast extract, 2% glucose) containing 0.6 M sorbitol was added, and the cells were incubated at 27°C for 3 h. After incubation, the cells were plated on SA medium containing nourseothricin (300 µg/mL) and incubated at 27°C for 3 days. Genomic DNA was extracted from the nourseothricin-resistant transformants, and deletion of the *atg1* gene was confirmed by PCR.

**Table 4.**
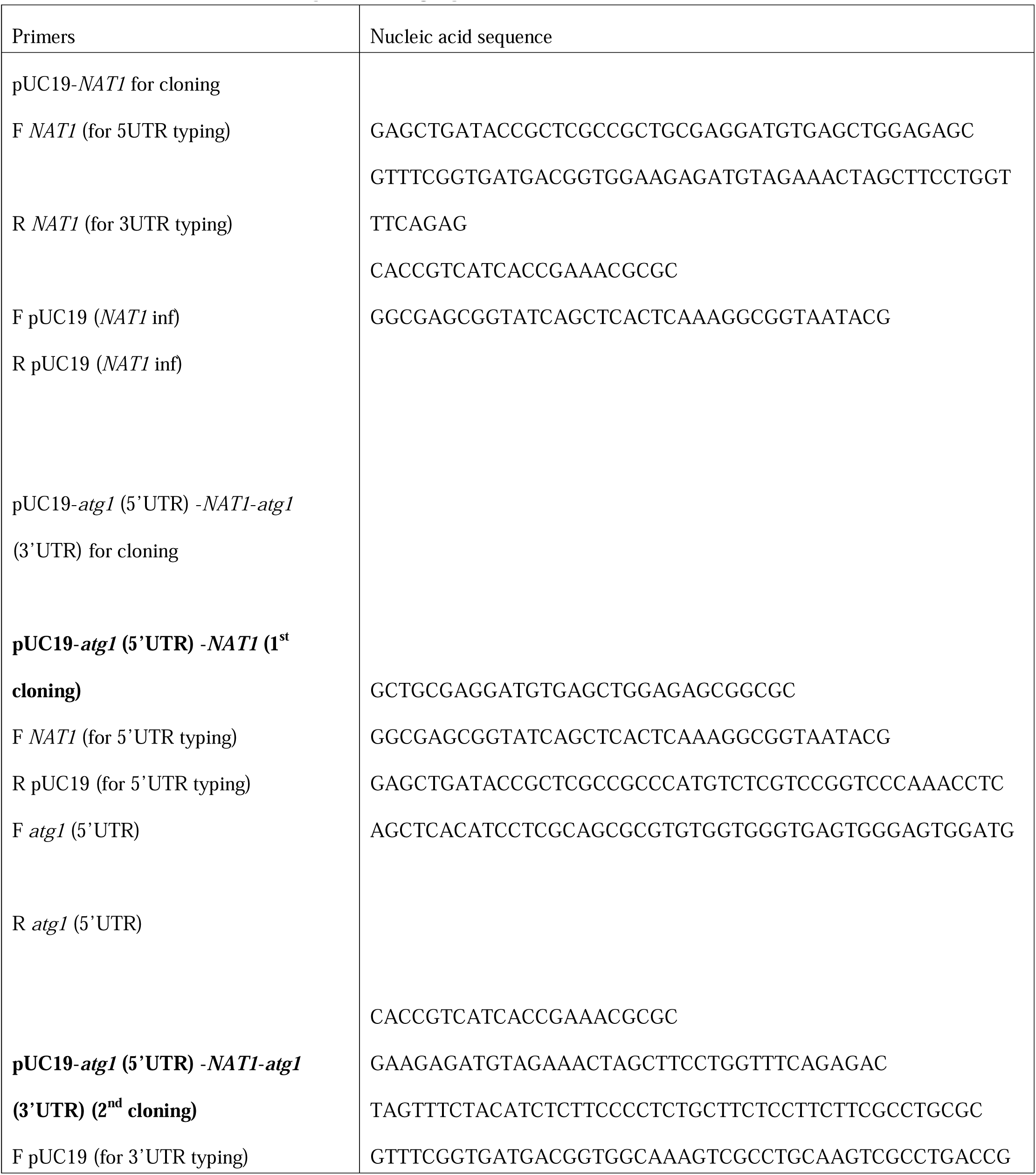
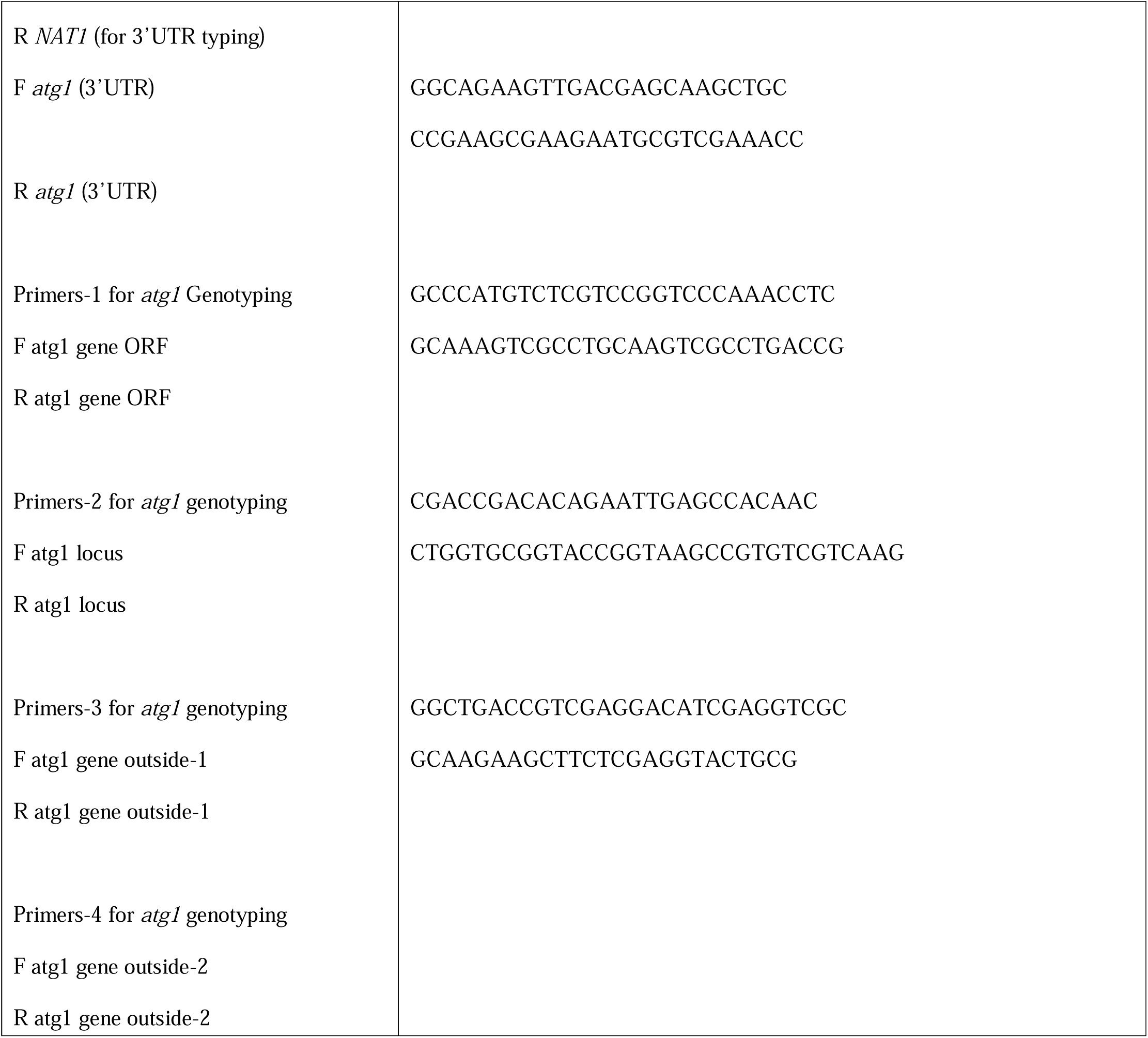
Primers used to generate *atg1* gene-deficient mutants.

### 2.6. Temperature sensitivity test

A temperature sensitivity test was performed as described previously with modifications (*20*,*21*). *T. asahii* strains were grown on SDA containing cefotaxime (100 μg/mL) at 27°C for 1 day. *T. asahii* cells were suspended in saline, and the absorbance at 630 nm was adjusted to 1. A 10-fold serial dilution of each suspension was prepared using saline. Five microliters of each dilution were spotted onto SDA and incubated at 27°C, 37°C, or 40°C for 48 h. Photographs of the agar plates were taken after incubation. For evaluation in liquid culture, Sabouraud dextrose broth (1% hipolypeptone, 4% dextrose) was used. *T. asahii* cells were suspended in saline and filtered through a 40-μm cell strainer. Absorbance at 630 nm was adjusted to 0.005. The suspensions were incubated at 27°C, 37°C, or 40°C for 96 h, and absorbance at 630 nm was measured every 24 h using a microplate reader (iMark™ microplate reader; Bio-Rad Laboratories Inc., Hercules, CA, USA).

### 2.7. Stress tolerance test

A drug sensitivity test was performed as previously reported with modifications (*20*,*21*). *T. asahii* was grown on SDA containing cefotaxime (100 µg/mL) at 27°C for 2 days. Cells were suspended in saline and filtered through a 40-µm cell strainer. Absorbance of the suspension was adjusted to 1.0 at 630 nm. A 10-fold serial dilution of each suspension was prepared using saline. Five microliters of each dilution were spotted onto SDA containing NaCl, KCl, Congo red, SDS, H O, tunicamycin, DTT, amphotericin B, or fluconazole. The plates were incubated at 27°C or 37°C for 24 to 72 h, and images of the agar plates were captured after incubation.

The minimum inhibitory concentration (MIC) was determined using the broth microdilution method according to CLSI guideline M27-A3, with a commercial antifungal susceptibility testing plate for yeasts (DP Eiken; Eiken Chemical Co., Ltd., Tokyo, Japan) (*21*). *T. asahii* was grown on SDA containing cefotaxime (100 μg/mL) at 27°C for 2 days. The cells were suspended in saline and adjusted to a turbidity equivalent to McFarland standard 1. The suspensions were then diluted 20-fold with saline, followed by a 100-fold dilution with RPMI 1640 medium (Thermo Fisher Scientific Inc., MA, USA) supplemented with 0.165 M 3-morpholinopropane-1-sulfonic acid (RPMI-MOPS, pH 7.0; FUJIFILM Wako Pure Chemical Corporation, Osaka, Japan). A 100-μL volume of the prepared cell suspension was added to each well of the plate containing one of the following eight antifungal agents: micafungin, caspofungin, amphotericin B, 5-fluorocytosine, fluconazole, itraconazole, voriconazole, or miconazole. The plates were incubated at 37°C for 48 h. MIC values were determined based on the results of two independent experiments.

### 2.8. Morphologic analysis of *T. asahii*

*T. asahii* strains were grown on SDA containing cefotaxime (100 µg/mL) at 27°C for 2 days. *T. asahii* cells were suspended in saline and filtered through a 40-µm cell strainer. Absorbance of the suspension was adjusted to 1.0 at 630 nm. A 20-µL aliquot of the suspension was added to 200 µL each of the Sabouraud dextrose broth (SB) and Synthetic Defined – Nitrogen (SD–N) medium. The cells were incubated at 37°C for 24 h. After incubation, culture samples were placed on microscope slides, covered with coverslips, and observed under a light microscope (BX51; OLYMPUS Corp., Tokyo, Japan). Photographs of the cells were randomly captured from multiple microscopic fields. The *T. asahii* morphology was classified as follows: yeast form, observed as single, spherical cells including budding yeasts; arthroconidia, observed as chains of three or more round or rectangular cells separated by septa; hyphal form, observed as elongated structures at least twice the length of yeast cells, including pseudohyphae, but lacking septa characteristic of arthroconidia (*21*,*24*). More than 400 cells were observed, and the number of each morphologic type (yeast, arthroconidia, hyphae) was counted.

### 2.9. Statistical analysis

All experiments were performed at least three times, and representative results are shown. For morphologic analysis, differences between groups were analyzed by Dunnett’s test. For silkworm infection assays, differences between groups were analyzed using the log-rank test based on Kaplan–Meier survival curves generated with EZR version 1.5.5 (https://www.jichi.ac.jp/saitama-sct/SaitamaHP.files/download.html). A *P*-value of < 0.05 was considered statistically significant.

## 3. Results

### 3.1. Identification and conservation of the Atg1 Protein in *T. asahii*

We sought to identify a gene encoding a homologous Atg1 protein in the *T. asahii* genome. In *C. neoformans*, *atg1* was identified based on the amino acid sequence of the *S. cerevisiae* Atg1 protein (*11*,*16*). *T. asahii* hypothetical protein A1Q1_02656 was identified as a candidate protein with 52% amino acid identity to *C. neoformans* Atg1 (Table 2). Next, we performed a homology analysis comparing the amino acid sequence of *T. asahii* hypothetical protein A1Q1_02656 with Atg1 proteins from representative fungi such as *S. cerevisiae*, *A. fumigatus,* and *C. albicans*. *T. asahii* hypothetical protein A1Q1_02656 showed 52%, 52%, 50%, and 47% amino acid identity with Atg1 proteins of *C. neoformans*, *A. fumigatus*, *C. albicans*, and *S. cerevisiae*, respectively (Figure 1A, Table 2). A phylogenetic tree generated from these amino acid sequences revealed that, among the fungi analyzed in this study, *T. asahii* hypothetical protein A1Q1_02656 was most closely related to *C. neoformans* Atg1 (Fig. 1B). Functional domain analysis revealed that the *T. asahii* hypothetical protein A1Q1_02656 also contained a PKc_like superfamily domain, a catalytic domain of protein kinases (Fig. 1C, Table 3). Therefore, the *T. asahii* hypothetical protein A1Q1_02656 was estimated to be the Atg1 protein in *T. asahii,* and the corresponding gene was designated *atg1*.

**Figure 1.**
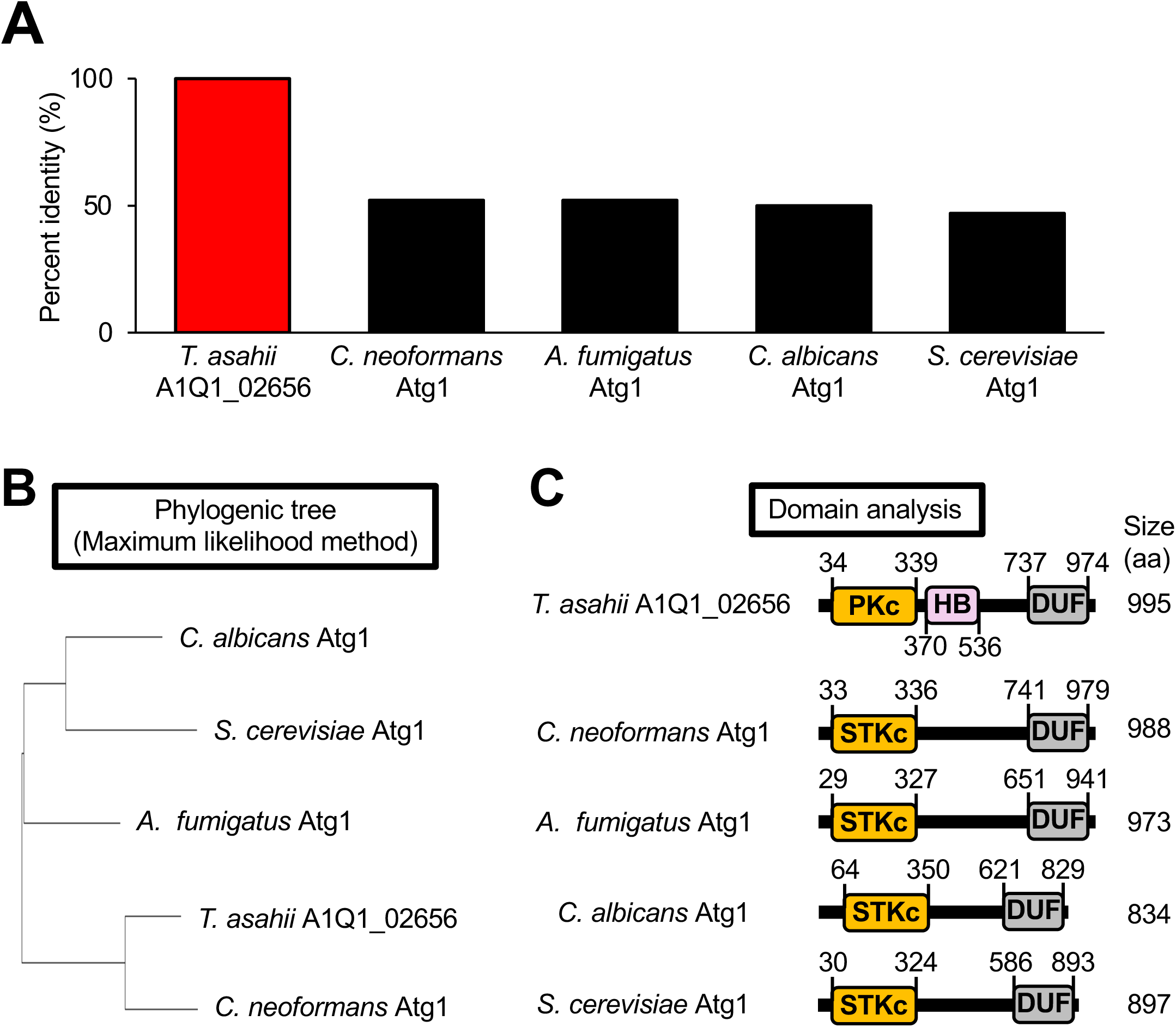
Conservation of *T. asahii* Atg1 among fungi. (**A**) Phylogenetic tree of Atg1 proteins from *T. asahii*, *C. neoformans*, *A. fumigatus*, *C. albicans*, and *S. cerevisiae*, constructed using the maximum likelihood method. (**B**) Phylogenetic tree of Atg1 proteins from *T. asahii*, *C. neoformans*, *A. fumigatus*, *C. albicans*, and *S. cerevisiae*, constructed using the maximum likelihood method. Phylogenetic analysis was performed using MEGA11 Software (https://www.ncbi.nlm.nih.gov/). (**C**) Schematic diagrams of domains present in Atg1 proteins. PKc: PKc_like superfamily domain. STKc: STKc_ATG1_ULK_like domain. HB: Herpes_BLLF1 superfamily domain. DUF: DUF3543. Functional domain prediction was performed using the National Center for Biotechnology Information (NCBI) (https://www.ncbi.nlm.nih.gov/).

### 3.2. Generation of *atg1* gene-deficient *T. asahii* mutants

The gene-targeting method used to generate gene-deficient *T. asahii* mutants was established previously (*20*,*25*). Using the method, we generated three *atg1* gene-deficient *T. asahii* mutants. To minimize the potential impact of secondary mutations, such as point mutations that may arise during mutant generation, three independent *atg1* gene-deficient strains of *T. asahii* were obtained from the same parental strain. The DNA fragment used for *atg1* gene deletion contained the *NAT1* gene, which confers resistance to nourseothricin (Fig. 2A). Homologous recombination replaced the *atg1* gene with the *NAT1* gene, thereby conferring resistance to nourseothricin (Fig. 2B). Transformation was performed by introducing the DNA fragment into the parental strain via electroporation, and transformants resistant to nourseothricin were obtained (Fig. 2B). Genomic DNA was extracted from the nourseothricin-resistant transformants, and *atg1* gene deficiency was confirmed by PCR (Fig. 2C, D). These results suggest that the *atg1* gene-deficient *T. asahii* mutants were generated.

**Figure 2.**
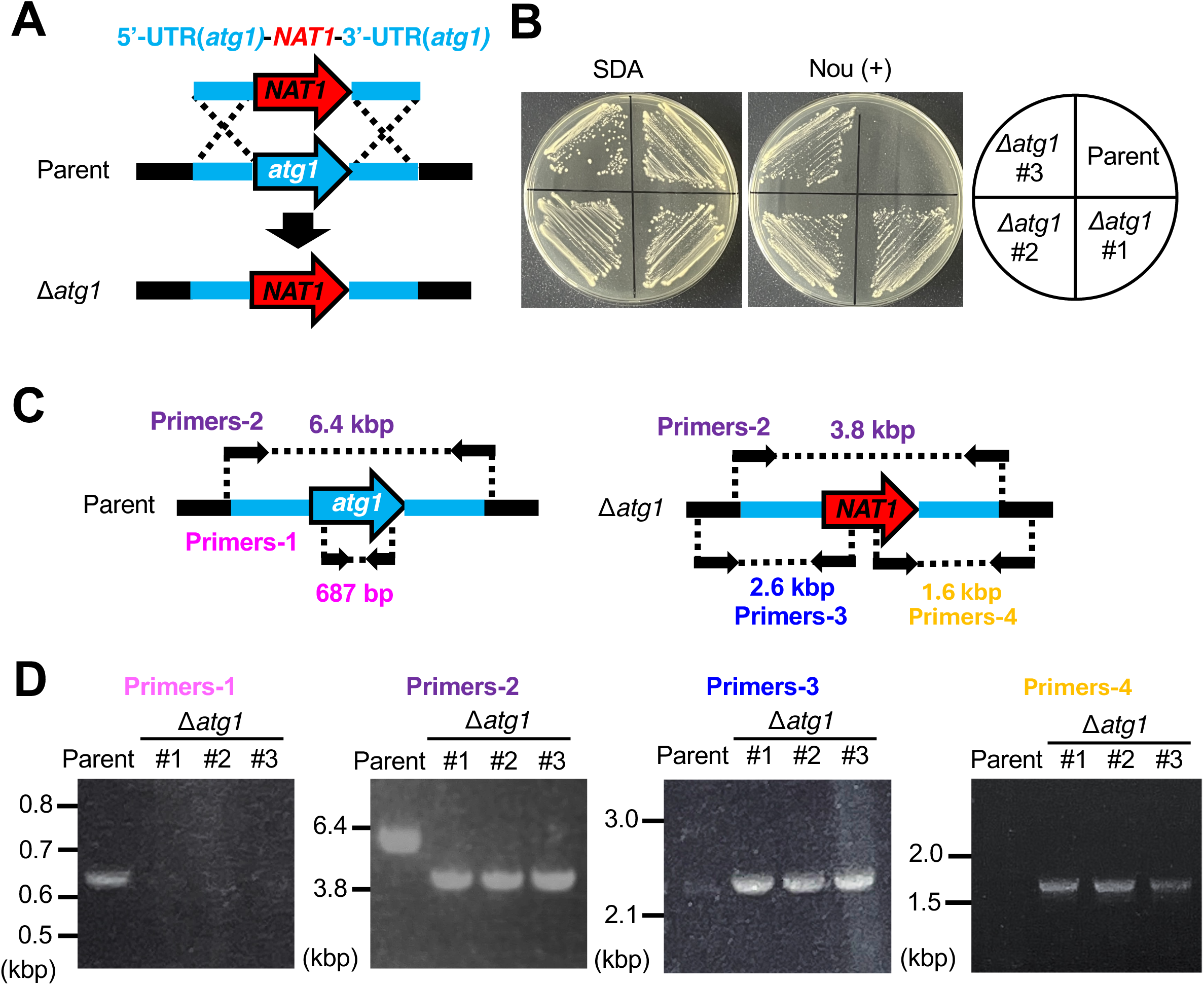
Generation of the *atg1* gene-deficient *T. asahii* mutants. (**A**) Schematic diagram of the gene recombination strategy used to construct the *atg1* gene-deficient *T. asahii* mutant (Δ*atg1*), along with the predicted genomic structure of the *atg1* gene-deficient mutant. (**B**) The parental strain (Parent) and the *atg1* gene-deficient mutants (Δ*atg1* #1-3) were spread on SDA or SDA containing nourseothricin (100 µg/mL; Nou (+)) and incubated at 27°C for 2 days. (**C**) Schematic illustration of primer positions and expected amplified DNA fragment sizes by PCR to verify the genomic structure of the *atg1* gene-deficient candidates using extracted genomic DNA. (**D**) DNA electrophoresis of PCR products to confirm the *atg1* gene deficiency (Δ*atg1*).

### 3.3. Effects of *atg1* gene deficiency on stress sensitivity

Next, we investigated whether growth of the *atg1* gene-deficient mutants was similar to that of the parental strain. Under temperatures of 27°C, 37°C, or 40°C, no significant differences in growth were observed between the parental strain and the *atg1* gene-deficient mutants (Fig. 3).

**Figure 3.**
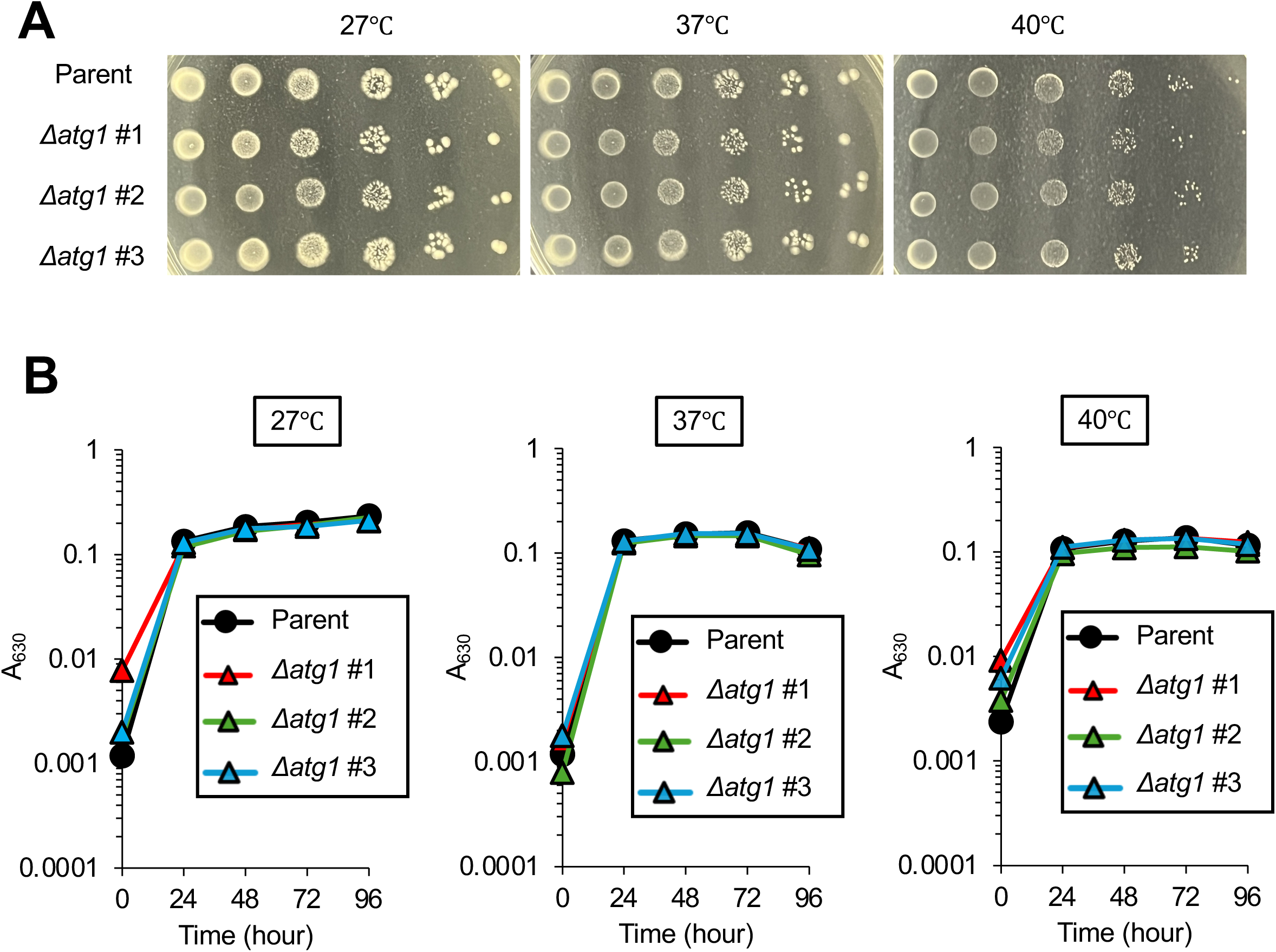
Growth of the *atg1* gene-deficient *T. asahii* mutants under high-temperature conditions. (**A**) The parental strain (Parent) and the *atg1* gene-deficient *T. asahii* mutants (Δ*atg1* #1-3) were grown on SDA at 27°C for 2 days. Cells were suspended in saline, and 10-fold serial dilutions were prepared. Each dilution (5 µl) was spotted onto SDA and incubated at 27°C, 37°C, or 40°C for 48 h. (**B**) The parental strain (Parent) and the *atg1* gene-deficient *T. asahii* mutants (Δ*atg1* #1-3) were grown in Sabouraud dextrose broth at 27°C, 37°C, or 40°C, and absorbance at 630 nm was measured. Data are presented as mean ± standard error of the mean (SEM).

In *C. neoformans*, mutants lacking autophagy-related genes exhibit sensitivity to oxidative and osmotic stress (*17*). To evaluate whether the *T. asahii atg1* gene is involved in osmotic stress, cell wall/membrane stress, endoplasmic reticulum stress, or oxidative stress, stress sensitivity assays were performed. The growth of the *atg1* gene-deficient mutants was not altered under the tested compounds (Fig. 4). The MICs of antifungal agents for the *atg1* gene-deficient strains were comparable to those for the parental strain (Table 5). These results suggest that the *atg1* gene is not involved in susceptibility to compound-induced stresses and antifungal agents in *T. asahii*.

**Figure 4.**
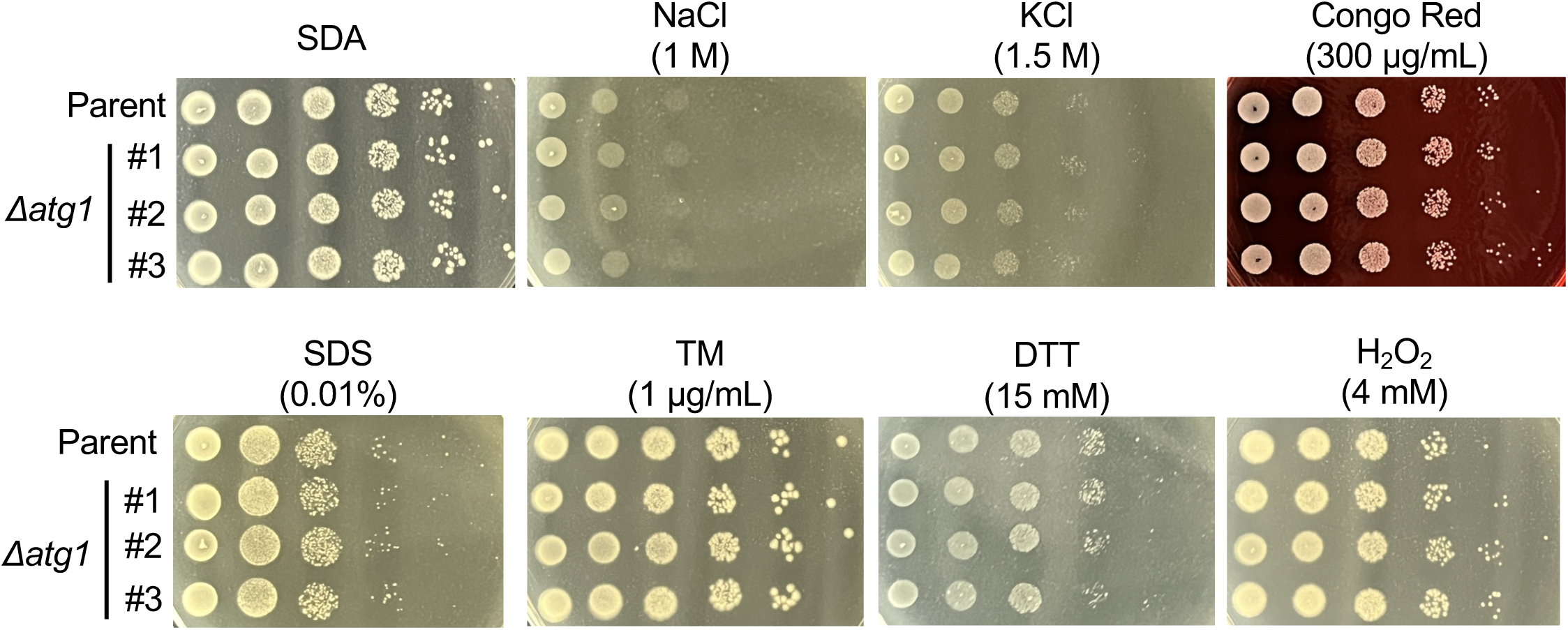
Growth of the *atg1* gene-deficient *T. asahii* mutants under stress-inducing conditions. The parental strain (Parent) and the *atg1* gene-deficient *T. asahii* mutants (Δ*atg1* #1-3) were cultured on SB at 27°C for 2 days. Cells were suspended in saline, and 10-fold serial dilutions were prepared. Each dilution (5 µL) was spotted onto SA medium containing NaCl (1 M), KCl (1.5 M), Congo red (300 µg/mL), SDS (0.01%), tunicamycin (TM, 1 µg/mL), dithiothreitol (DTT, 15 mM), or hydrogen peroxide (H O, 4 mM), and incubated at 27°C for 48 h.

**Table 5.** MIC values of *atg1* gene-deficient mutants against antifungal agents.

| MIC (μg/mL) | Parent | *Δatg1* #1 | *Δatg1* #2 | *Δatg1* #3 |
| --- | --- | --- | --- | --- |
| MCFG | >16 | >16 | >16 | >16 |
| CPFG | 4 | 4 | 4 | 4 |
| AMPH-B | 0.5 | 0.5 | 0.5 | 0.5 |
| 5-FC | >64 | >64 | >64 | >64 |
| FLCZ | 1 | 1 | 1 | 1 |
| ITCZ | 0.5 | 0.5 | 0.5 | 0.5 |
| VRCZ | 0.12 | 0.12 | 0.12 | 0.12 |
| MCZ | 0.5 | 0.5 | 0.5 | 0.5 |

The *T. asahii* parental strain (Parent) and three independent *atg1* gene-deficient mutant *T. asahii* strains (*Δatg1* #1-3) were cultured in Sabouraud medium for 2 days at 27 °C. *T. asahii* cells were suspended in saline, and cell suspensions were prepared to a turbidity of 1 according to the McFarland turbidity method. Each bacterial solution was diluted 20-fold in saline and then 100-fold in RPMI MOPS. The prepared fungal solutions were then diluted with eight antifungal agents (Micafungin [MCFG], caspofungin [CPFG], amphotericin B [AMPH-B], 5-fluorocytosine [5-FC], fluconazole [FLCZ], itraconazole [ITCZ], voriconazole [VRCZ], miconazole [MCZ]) and inoculated on plates at 37°C for 48 h. The minimum inhibitory concentration (MIC) was determined from the results of two independent experiments.

### 3.4. Role of the *atg1* gene in *T. asahii* morphology

*T. asahii* exhibits dimorphism, alternating between yeast forms and filamentous/arthroconidial forms (*1*,*24*). In *C. albicans*, a dimorphic fungus, hyphal formation is induced by various nutritional conditions, such as growth at 37°C and exposure to GlcNAc (*26*). To observe the morphologic changes of *T. asahii* yeast cells under various nutritional conditions, we developed a method to collect *T. asahii* yeast cells. More than 90% of *T. asahii* cells in suspension filtered through a 40-µm filter were yeast cells (Fig. 5A, B). Using the *T. asahii* yeast cells, we tested the growth of the cells in various culture media, such as SB and SD-N. SB is a commonly used medium for culturing *T. asahii*, whereas SD-N is a nutrient-limited medium with a restricted nitrogen source (*24*). The absorbance of *T. asahii* culture medium at 630 nm was increased in the order SB and SD-N (Fig. 5C). The ratio of hyphae was increased in SB by incubating at 37°C for 24 h (Fig. 5D, E). The ratio of arthroconidia was also increased by incubation in SB and SD-N at 37°C for 24 h (Fig. 5D, E). We investigated whether the *atg1* gene contributes to the morphologic changes of *T. asahii* in various culture media. The *atg1* gene-deficient mutants exhibited a higher proportion of hyphal cells in SB (Fig. 6). No significant morphologic differences were observed, however, between the parental strain and the *atg1*-deficient *T. asahii* mutants in SD-N (Fig. 6). These results suggest that the *atg1* gene negatively regulates hyphal formation in *T. asahii* in a nutrient-dependent manner.

**Figure 5.**
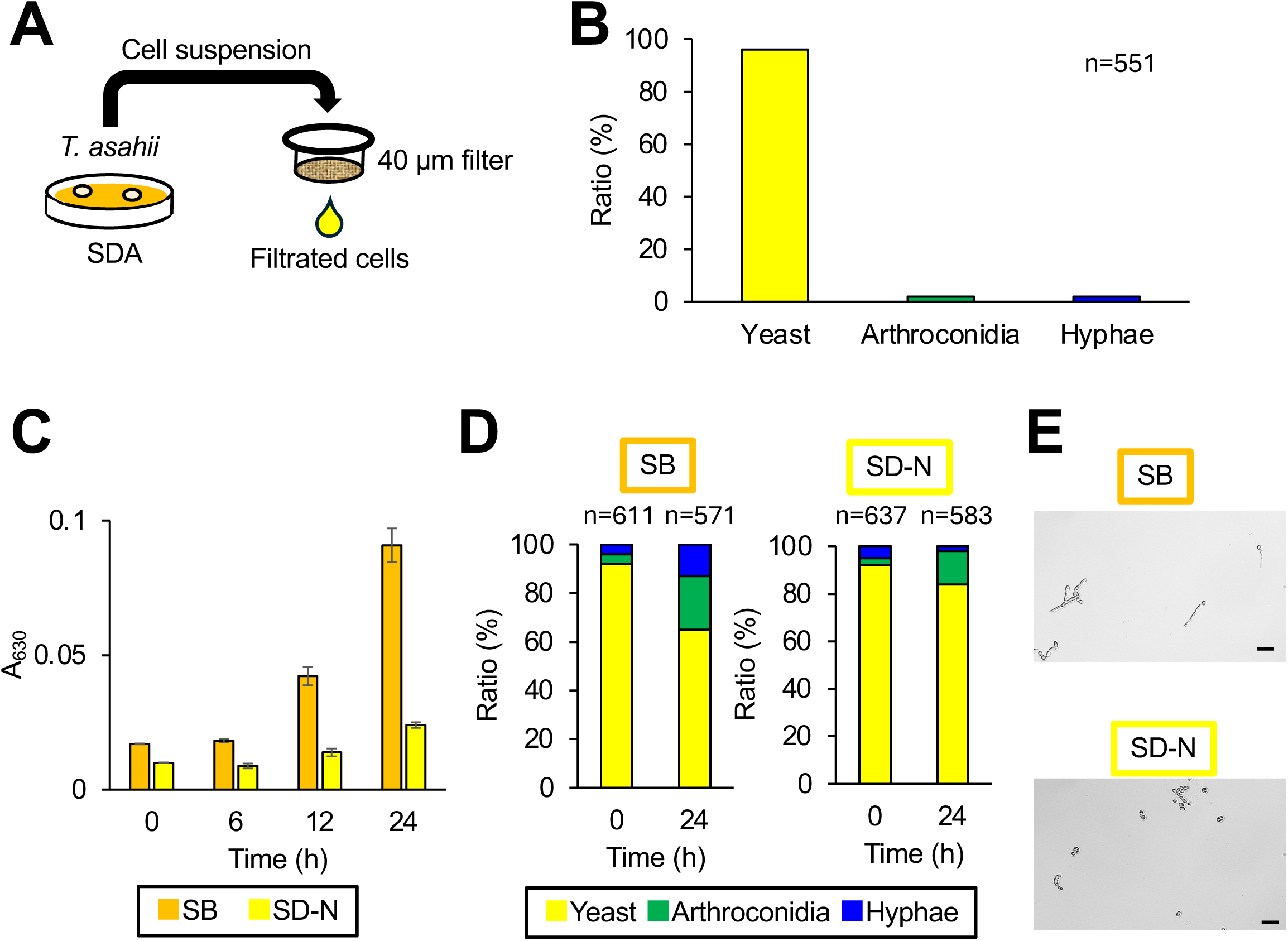
Effects of culture conditions on *T. asahii* morphology. (**A**) Method for preparing the yeast form of *T. asahii*. The parental *T. asahii* strain (Parent) was cultured on SDA at 27°C for 2 days. *T. asahii* cells were suspended in saline and filtered through a 40-µm cell strainer. (**B**) The ratio of each cell shape in the filtered *T. asahii* cells was measured. (**C**) Absorbance of the cell suspension at 630 nm was adjusted to 0.5, and 5 µL of the filtered cell suspension was added to 200 µL of Sabouraud dextrose medium or SD-N liquid medium. Cultures were incubated at 27°C for 24 h, and absorbance at 630 nm was measured for 24 h. (**D-F**) Absorbance of the filtered cell suspension at 630 nm was adjusted to 1.0, and 20 µL of the suspension was added to 200 µL of Sabouraud dextrose broth (SB) or synthetic defined – nitrogen (SD–N) medium, followed by incubation at 27°C for 24 h. Cells were then observed under a light microscope. Scale bars represent 40 µm. Images were captured randomly from multiple microscopic fields. Cell numbers in three morphologic categories: yeast, arthroconidia, and hyphae, were quantified. (**D**) Cell numbers in three morphologic categories were measured at 0 or 24 h after incubation at 37°C. (**E**) Photographs of cells after incubation at 37°C for 24 h were obtained under microscopic fields. *n* represents the total number of microscopic fields analyzed from two independent experiments.

**Figure 6.**
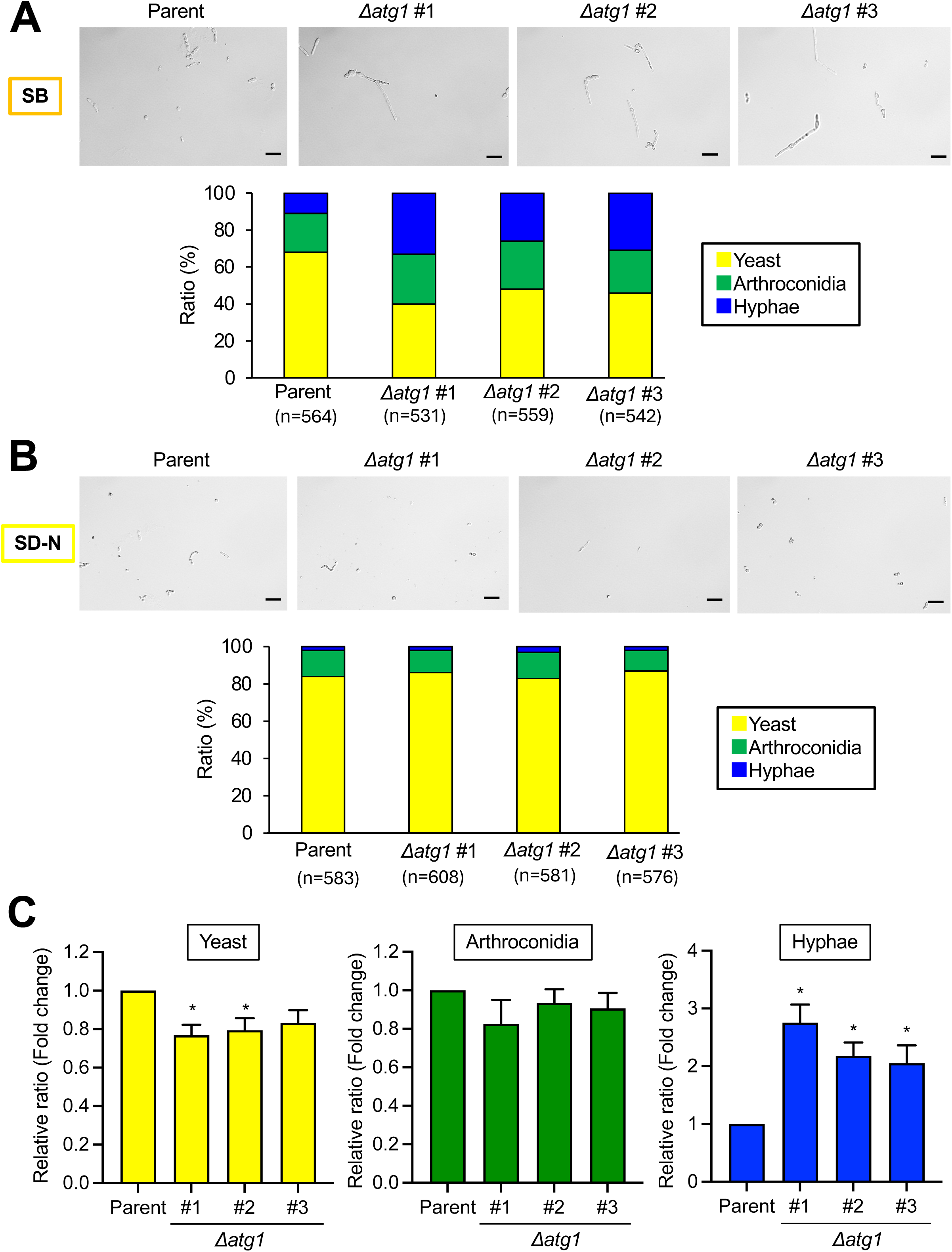
Effect of *atg1* gene deficiency on *T. asahii* morphology. The parental *T. asahii* strain (Parent) and three *atg1* gene-deficient mutants (Δ*atg1* #1-3) were grown on SDA at 27°C for 2 days. Cells were suspended in saline and filtered through a 40-µm cell strainer. Absorbance at 630 nm was adjusted to 1. A 20-µL aliquot of each suspension was added to (**A**) Sabouraud dextrose broth (SB), or (**B**) synthetic defined – nitrogen (SD–N) medium (200 µL). The samples were incubated at 37°C for 24 h. After incubation, the *T. asahii* cells were dropped onto microscope slides, covered with coverslips, and observed under a light microscope. Upper panels show representative microscopic images. Scale bars represent 20 µm. Lower panels show quantification of the three morphologic forms—yeast, arthroconidia, and hyphae—based on randomly selected microscopic fields. The total number of cells was obtained from two independent experiments. (**C**) The relative ratio of the three morphologic forms were determined compared to the ratio of parental strain (Parent) under a culture condition with SB at 37°C for 24 h. The data of the relative ratio were obtained from seven independent experiments. Error bars indicate the SEM of the means (n = 7). Statistical differences between parental strain and Δ*atg1* #1-3 were analyzed by Dunnett’s test. (*: *P* < 0.05).

### 3.5. Virulence of the *atg1* gene-deficient mutants in a silkworm infection model

In *C. neoformans*, *atg1 g*ene deficiency did not affect its virulence in a *Galleria mellonella* infection model (*17*). We tested whether the *atg1* gene is required for *T. asahii* virulence in a silkworm infection model. The survival time of silkworms infected with the *atg1*-deficient mutants was not significantly different from that of silkworms infected with the parental strain (Fig. 7). The result suggests that the *atg1* gene does not contribute to *T. asahii* virulence in the silkworm model.

**Figure 7.**
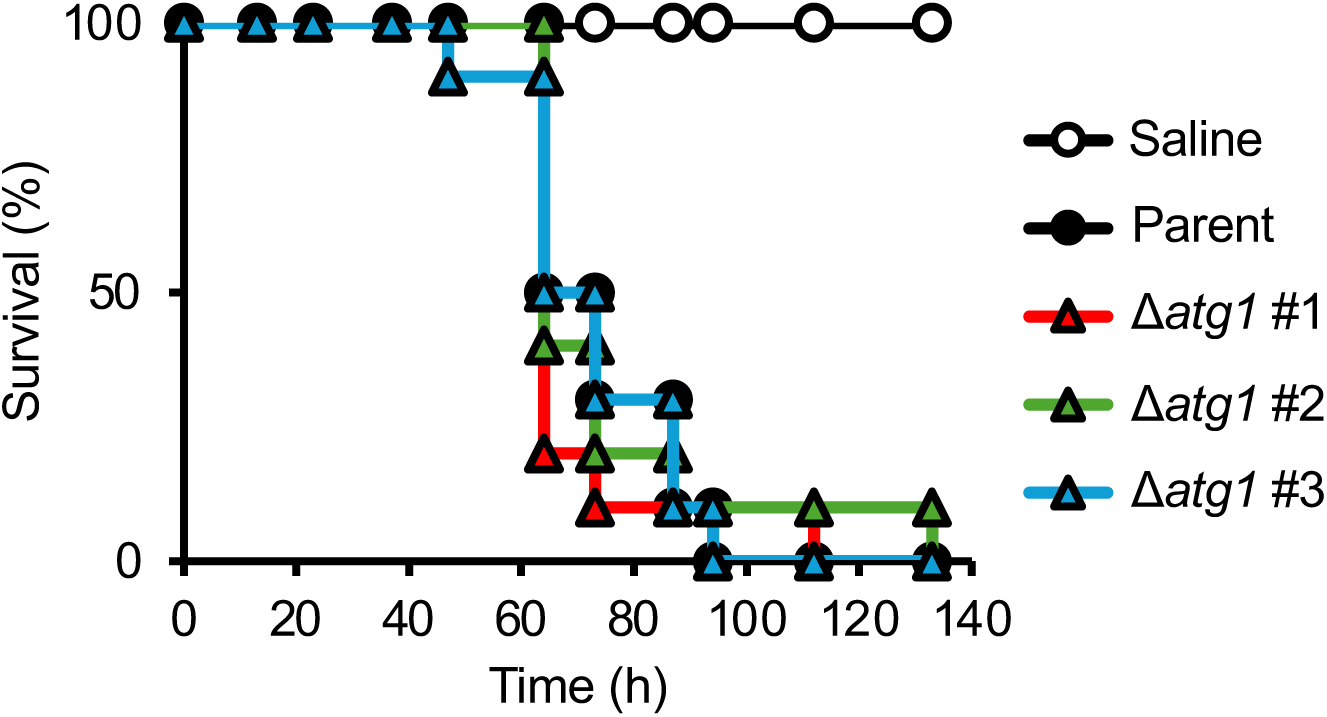
Virulence of *T. asahii atg1* gene-deficient mutants in a silkworm infection model. The parental strain (Parent; 530 cells/larva) and three independent *atg1* gene-deficient *T. asahii* mutants (Δ*atg1* #1-3; 790 - 950 cells/larva) were injected into the hemocoel of silkworms, and survival was monitored at 37°C for 96 h. Statistical significance between the parental and *atg1*-deficient groups was determined by the log-rank test based on Kaplan–Meier survival curves. Ten silkworms per group were used.

## 4. Discussion

The findings of the present study demonstrated that the *atg1* gene is involved in morphologic transitions of *T. asahii* under culture conditions using SB, a medium commonly used for fungi. On the other hand, *atg1* gene deficiency did not affect growth under various stress conditions or virulence of *T. asahii* in the silkworm model. These findings suggest that Atg1 might play a role in regulating hyphal formation under specific culture conditions but is not involved in stress resistance or virulence in *T. asahii*.

*T. asahii* Atg1 protein was estimated based on the amino acid sequence of other fungal Atg1 proteins and their domain analysis. *T. asahii* hypothetical protein A1Q1_02656 has high sequence similarity to the Atg1 proteins of other representative fungi, including *S. cerevisiae*, *A. fumigatus*, and *C. albicans*. Domain analysis revealed that *T. asahii* hypothetical protein A1Q1_02656 harbors a Pkc_like superfamily domain—a catalytic domain of protein kinases conserved among the analyzed fungi. In *S. cerevisiae*, Atg1 kinase activity contributes to non-selective autophagy (*9*). Therefore, we determined that *T. asahii* hypothetical protein A1Q1_02656 is the Atg1 protein of *T. asahii*.

*T. asahii atg1* gene deficiency does not significantly affect the stress response and antifungal drug resistance of *T. asahii*. We generated *atg1* gene-deficient *T. asahii* mutants and analyzed their phenotypes. When generating gene-deficient mutants, unintended secondary mutations may be introduced in addition to the targeted gene disruption. To minimize the possibility of this rare occurrence, we generated three independent *T. asahii atg1* gene-deficient mutants in this study and examined whether they exhibited consistent phenotypes. The *atg1* gene deficiency in *T. asahii* did not affect sensitivity to high temperature, NaCl, KCl, tunicamycin, DTT, H_2_O_2_, or antifungal drugs. In *C. neoformans*, the *atg1* gene-deficient mutant also did not exhibit high-temperature and H_2_O_2_-sensitive phenotypes (*16*,*17*). Therefore, the *atg1* gene is not involved in high temperature and oxidative stress tolerance in *T. asahii* or *C. neoformans*. On the other hand, in *C. neoformans*, the growth of the *atg1* gene-deficient mutant was delayed by NaCl (*17*). The *atg1* genes in *T. asahii* and *C. neoforman*s might differ in their tolerance to NaCl. The calcineurin and Hog1 pathways are involved in high-temperature tolerance in *T. asahii* (*20*,*21*). We therefore concluded that the *T. asahii* Atg1 is regulated in a calcineurin and Hog1 pathway-independent manner.

The *atg1* gene contributes to the morphologic changes of *T. asahii*. In SB, the ratio of hyphae was increased by *atg1* gene deficiency after 24-h incubation. On the other hand, in SD-N, a nitrogen-limited medium, the morphology in the *atg1* gene-deficient mutants was not different from that in the parental strain. In fungi such as *S. cerevisiae*, *C. neoformans*, and *C. albicans*, Atg1 is involved in nitrogen limitation-induced autophagy and required for survival under nitrogen-limited conditions (*27*). In the present study, we revealed the role of Atg1 in the morphologic changes of *T. asahii* under culture conditions using SB. Further studies are needed to elucidate the intracellular mechanisms regulated by Atg1 under culture conditions using SB. Calcineurin deficiency in *T. asahii* increases the proportion of the yeast form (*20*). In contrast, Hog1 deficiency increases the proportion of hyphal forms (*21*,*28*). The involvement of the calcineurin and Hog1 pathways in *atg1* gene-related morphological transformation in *T. asahii* remain to be elucidated.

The *T. asahii atg1* gene is not involved in *T. asahii* virulence in the silkworm infection model. In *C. neoformans*, Atg1 does not influence virulence in the insect model *Galleria mellonella* (*17*). Therefore, we assume that Atg1 does not have a pivotal role in *T. asahii* and *C. neoformans* virulence in insect models. The Hog1 pathway contributes to morphological transformation, stress tolerance, and virulence against silkworms in *T. asahii* (*21*,*25*,*28*). We assumed that the increased rate of hyphal formation observed in *atg1* gene-deficient mutants appears insufficient to alter virulence against silkworms.

A limitation of this study is that the effects of *atg1* gene-deficiency of *T. asahii* on autophagy were not evaluated. Autophagy validation methods have been established in several fungi using fungal cells expressing GFP-Atg8 (*29*). The development of a detection system for autophagy in *T. asahii* using cells expressing GFP-Atg8–expressing cells remains a future challenge.

In conclusion, the *atg1* gene contributes to the morphologic changes of *T. asahii* under culture conditions using SB. Hyphal formation of *T. asahii* might be partially regulated by Atg1-dependent cellular events.

## Acknowledgements

We thank Renta Endo, Momoka Yagi, Haruka Ogino, Momoka Matsumura, and Tomoya Sanbongi (Meiji Pharmaceutical University) for technical assistance in rearing the silkworms. We also thank SciTechEdit International LLC (Highlands Ranch, CO, USA) for editorial support.

